# Differences in memory for what, where, and when components of recently-formed episodes

**DOI:** 10.1101/2020.01.09.901033

**Authors:** John J. Sakon, Roozbeh Kiani

## Abstract

An integral feature of human memory is the ability to recall past events. What distinguishes such episodic memory from associative and semantic memories is the joint encoding and retrieval of “what,” “where,” and “when” (WWW) of events. Here, we investigated whether the WWW components of episodes are retrieved with equal fidelity. Using a novel task where human participants were probed on the WWW components of a recently-viewed synthetic movie, we found fundamental differences in mnemonic accuracy between these components. The memory of “when” had the lowest accuracy and was most severely influenced by primacy and recency. Further, the memory of “when” and “where” were most susceptible to interference due to changes in memory load. These findings suggest that episodes are not stored and retrieved as a coherent whole. Rather, memory components preserve a degree of independence, suggesting that remembering coherent episodes is an active reconstruction process.

---

Episodic memories store past events that shape how we remember our lives. Each episodic engram unites disparate streams from our sensory cortices into a combined representation^1^. At minimum, episodic memories bind “what”, “where”, and “when” (WWW) components into these engrams^2,3^. Therefore, studies in animals^4-8^ and humans^1,2,9-11^ have focused on these key components when probing the presence of—and brain structures responsible for—episodic memory.

However, little is known about the strength of association and interactions between the WWW components of episodic memory^12-14^. Put more simply: are the what, where and when components of episodic memory encoded and recalled with equal fidelity? One possibility is that an episodic engram is a holistic representation in which the WWW components are inseparably intertwined. In this case, encoding and retrieval of each component should invariably be at the same level as the other components. Alternatively, the WWW components may be stored separately, and re-joined post-hoc during retrieval to synthesize a coherent memory of a past event. Between these two extreme hypotheses, there is a spectrum in which the what, where, and when components of an event are encapsulated with various degrees of separability in an engram.

A handful of studies have investigated this separability between the components of episodes^1,12-19^. An early study that tasked humans to remember series of consonants that differed in either spatial (where) or temporal (when) order found limited differences in recall accuracy once phonemic coding (saying the letters to remember them) was accounted for^12^. A more recent study attempting to probe the separability of spatial vs. temporal memory utilized objects with little resemblance to real-world objects to avoid such phonemic confounds^13^. Following serial presentation of objects during a study phase, spatial and temporal memory were probed by altering the location and/or order of the objects during the test phase. A change in temporal order affected spatial memory accuracy when subjects were asked to focus on location memory, while a change in spatial relations did not affect temporal memory accuracy when subjects were asked to focus on temporal order^13^. In addition, a series of studies where subjects saw or heard a sequence of items found better memory of temporal order of the items than their presented location, but only when explicitly told to pay attention to serial order^14-16^. Another study found better mnemonic accuracy for object or location memory when the task was related to that component^19^.

However, to the best of our knowledge no study has simultaneously probed the what, where, and when components of episodes in a single task design. Further, as summarized in the last paragraph, studies that have contrasted the interaction between two of the WWW components typically only reported differences in accuracy when subjects were tasked to focus on one of the components^13,15-17,19^. We overcome these limitations with a novel task that tests the memory for all three WWW components of episodes without instructing subjects to pay attention to any particular component. In this task, which we call TRANSFER (**TRAN**sient **S**napshots **F**rom **E**pisodic **R**ecall), subjects first view a movie and are asked to memorize it (encoding phase) (Fig. 1). The movie contains a series of shapes or “features” that sequentially appear and disappear in different locations. Subjects are then probed on the separate WWW components of their memory (retrieval phase) by classifying still images as matches or mismatches from movie content.

**Figure 1.**
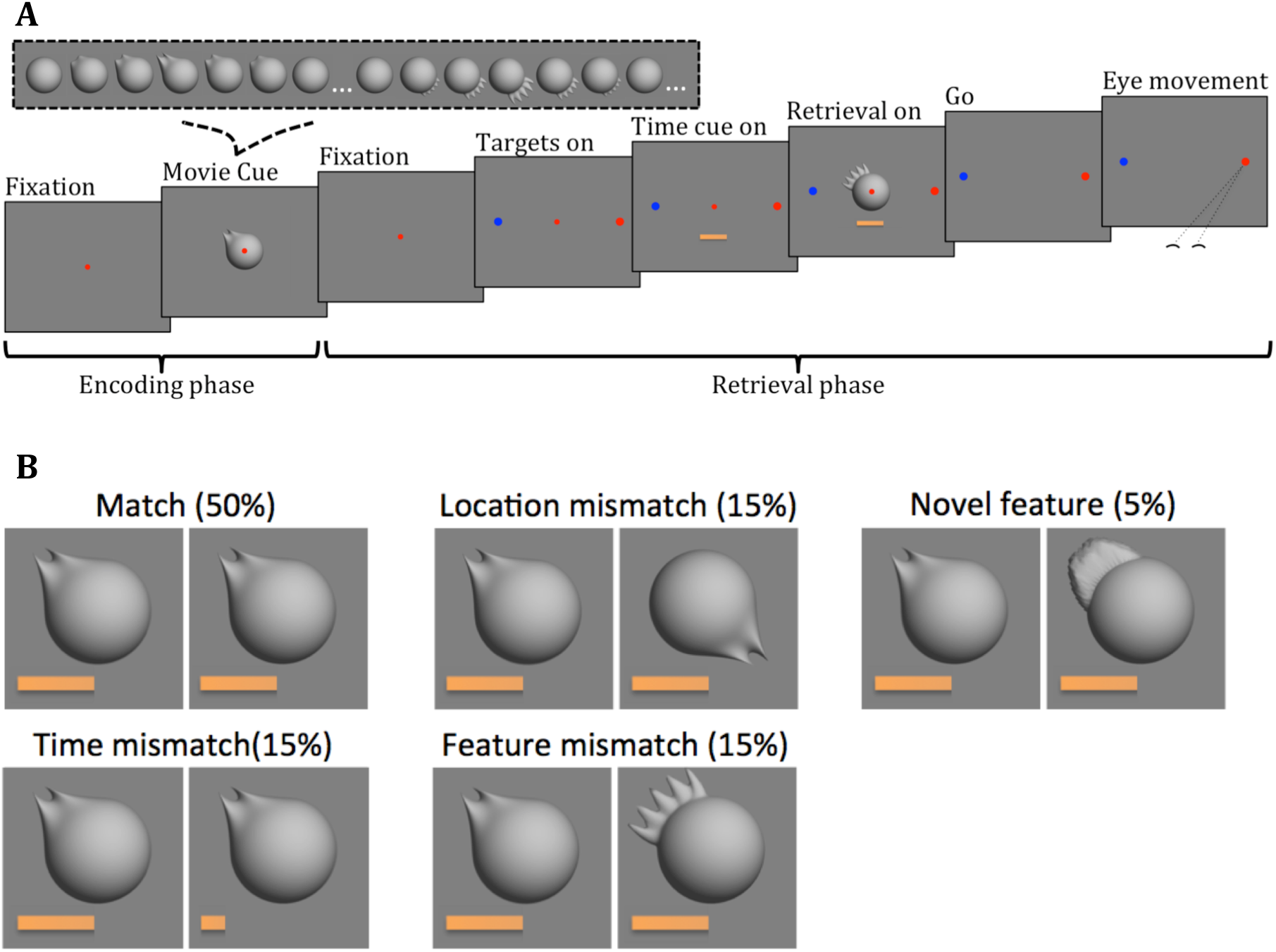
Outline of the TRANSFER task. **A)** Task design. Each block started with an encoding phase in which a movie with 3-6 features was shown (inset: 2 example features, with white ellipses indicating 2 s inter-feature intervals). The movie was followed by a retrieval phase that included 1-3 trials. In each retrieval trial subjects judged whether a feature and its location occurred in the movie at a cued time. **B)** Example of the 5 possible retrieval trial types. The retrieval trials in each block were chosen randomly according to the percentages for each type.

Our null hypothesis was that each of the WWW components of episodes were stored and retrieved with equal fidelity. However, we found evidence of differences between the three components, with trials probing “when” memory showing both the lowest accuracy and the most dramatic decreases in accuracy due to the serial order of features in the movie. In addition, when we increased the mnemonic load on the “what” component by showing two similar features at different times in the movie, the accuracy of “when” and “where” memory was strongly affected. Based on our results, we conclude that episodes are not stored and retrieved as a coherent whole, but instead tie the WWW components together via an engram in which certain components can be remembered with more or less fidelity.

## Results

To study how strongly different components of episodes were associated in memory, we developed the TRANSFER task, in which subjects observed a movie and after a short delay recalled events that occurred at particular times of the movie (Fig. 1A). To ensure equal complexity and salience of the events throughout the movie, we created synthetic movies in which a series of 3D features sequentially appeared and disappeared on the surface of a central sphere (Fig. 1A, inset). The features had an equal number of vertices, appeared in clearly discrete locations (four locations, 45°, 135°, 225°, or 315° around the circumference), were comfortably separated in time (2 s from end of one feature protrusion until the start of the next), and were easily distinguishable from each other (except in a manipulation condition explained below). In each block of the task, subjects first observed a movie with 3-6 features (encoding phase), and then were probed about features shown at particular times in the movie (retrieval phase). Between one and three retrieval trials occurred in the retrieval phase of each block depending on the length of the movie (see Methods for details). In each retrieval trial, a progress bar cued a time in the movie that subjects had to recall. Then, a still image of a feature at a particular location on the sphere was shown. Subjects indicated with a saccadic eye movement to one of two targets whether the shape and location of the feature in the still image matched or mismatched the cued time in the movie.

Mismatches could be due to (i) a movie feature that did not match the cued time but shown in the location matching the cued time (**feature-mismatch**), (ii) a movie feature matching the cued time but shown in a mismatching location (**location-mismatch**), (iii) a feature in its original location from a different time in the movie (**time-mismatch**), or (iv) a **novel-feature** not shown in the movie (Figure 1B). To perform well, subjects had to memorize episodes that encapsulated the location, shape, and time of the features in the movie. The novel-feature trials tested subjects’ recognition memory, but the other three trial types tested the what, where, and when (WWW) components of the episode. This aspect of the TRANSFER task is unique from previous work in that each of the WWW components of episodes can be separately probed using the feature-mismatch, location-mismatch, and time-mismatch trials, respectively. Importantly, subjects were given no instruction to focus on any particular component of the movie episodes and the three trial types were tested at equal proportions, making each component of the episodes equally important for achieving a high performance.

A key design principle of the TRANSFER task was to make the sensory signals of the shape, location, and presentation time of features strong and easily distinguishable, leaving no uncertainty about what, where, and when each feature was shown as subjects watched the movie. To put this another way, it would be trivial for subjects to achieve very high accuracy if we separately tested the three components using match-mismatch tasks that 1) probed only feature identity (“what”) of two features side-by-side in the same movie, 2) probed only location (“where”) of the same two features side-by-side in the same movie, and 3) probed only temporal order (“when”) by showing a movie with two features in the same location one after another^20^. Therefore, lower than close-to-perfect accuracies as well as differences in accuracy for each mismatch trial type were shaped by the memory mechanisms that underlie the encoding and retrieval of multi-feature episodes in the TRANSFER task.

If the WWW components of episodes were inseparable in memory we would expect similar accuracies for different mismatch trial types. We tested this hypothesis with two series of analyses: direct comparison of accuracy between trial types, and determining whether the accuracies for each trial type were equally influenced by three common mnemonic phenomena: primacy, recency, and interference.

### Different components of episodes have unequal retrieval accuracy

The retrieval accuracies were systematically different across trial types (Fig. 2; average pairwise difference, 5.1±3.5%). These differences were highly significant (p < 10^−8^, likelihood ratio test, Eq. 2 vs. Eq. 3), remained if we removed novel-feature trials (p = 2.4 × 10^−7^, likelihood ratio test, Eq. 2 vs. Eq. 3), and were consistent across subjects (Fig. S1). The lowest accuracy was for time-mismatch trials, which was significantly lower than match trials (*β*_1_ =0.30 ± 0.060, p = 1.3×10^−6^; Eq. 2, FWER-corrected), location-mismatch trials (*β*_2_ =0.23 ± 0.082, p = 0.024; Eq. 2, FWER-corrected), feature-mismatch trials (*β*_3_ =0.43 ± 0.088, p = 3.3×10^−6^; Eq. 2, FWER-corrected), and novel-feature trials (*β*_4_=1.4 ± 0.17, p < 10^−8^; Eq. 2, FWER-corrected). The highest accuracy was for novel-feature trials, which were significantly higher than all other mismatch trials (*β*_1_ =-1.0 ± 0.17, p < 10^−8^; *β*_2_ = −1.4 ± 0.17, p < 10^−8^; *β*_3_ = −1.1 ± 0.17, p < 10^−8^; *β*_4_ = −0.92 ± 0.18, p = 8.8×10^−7^; Eq. 4, all FWER-corrected).

**Figure 2.**
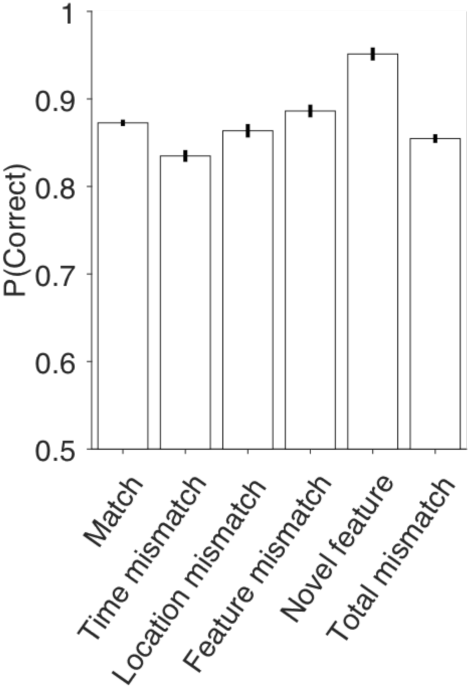
Different accuracies for what, where, and when components of episodes. The five bars on the left show the accuracy for the different trial types. The rightmost bar shows the overall accuracy of the four mismatch trial types. Data are pooled across 10 subjects. For single subject results see Fig. S1. Only non-morph blocks are used to make the figure (see Methods). Error bars are standard error of mean (SEM).

The higher accuracy for the novel-feature trials is expected because a correct answer in these trials can be achieved from recognition memory^21^, making them less likely to rely on recall mechanisms. However, the differential accuracy for the other mismatch trial types provides initial evidence that the what, where, and when components of episodes are not packaged in an inseparable fashion.

### Primacy and recency effects differ across components of episodes

When subjects are tasked to remember serially presented items, a common finding is that memory of items presented early (primacy) or late (recency) in the order are remembered better than items presented in the middle^22^. We investigated the strength of primacy and recency in our data by plotting the accuracy on retrieval trials as a function of cued time in the movie.

Primacy and recency effects were both present in our experiment. Figure 3A shows changes of retrieval accuracy as a function of cued time from the beginning (Fig. 3A left) or end (Fig. 3A right) of the movies. Aggregated across all movie lengths and trial types, accuracy was systematically higher for the earliest (*β*_1_ = 0.21 ± 0.014, p < 10^−8^; Eq. 5) and latest (*β*_2_ = 0.32 ± 0.044, p < 10^−8^; Eq. 5) features in the movies. These effects were not due to variable movie lengths across blocks as they were present for each movie length over 3 features (Fig. S3), but they were strongest for the longest movies, as expected (results from blocks with six-feature movies are depicted in Fig. 3B). Also, the primacy and recency effects were strongly present in the data from all subjects: 10/10 subjects showed better accuracy for the first cued feature compared to the second (p=0.002, exact binomial test), and for the last feature compared to the second-from-last (p=0.002, exact binomial test; Fig. S2A-B). Primacy and recency slopes were also significant when fit to individual subject’s behavior (primacy: p=0.011; recency: p=0.0010; one-way sign test, Eq. 5).

**Figure 3.**
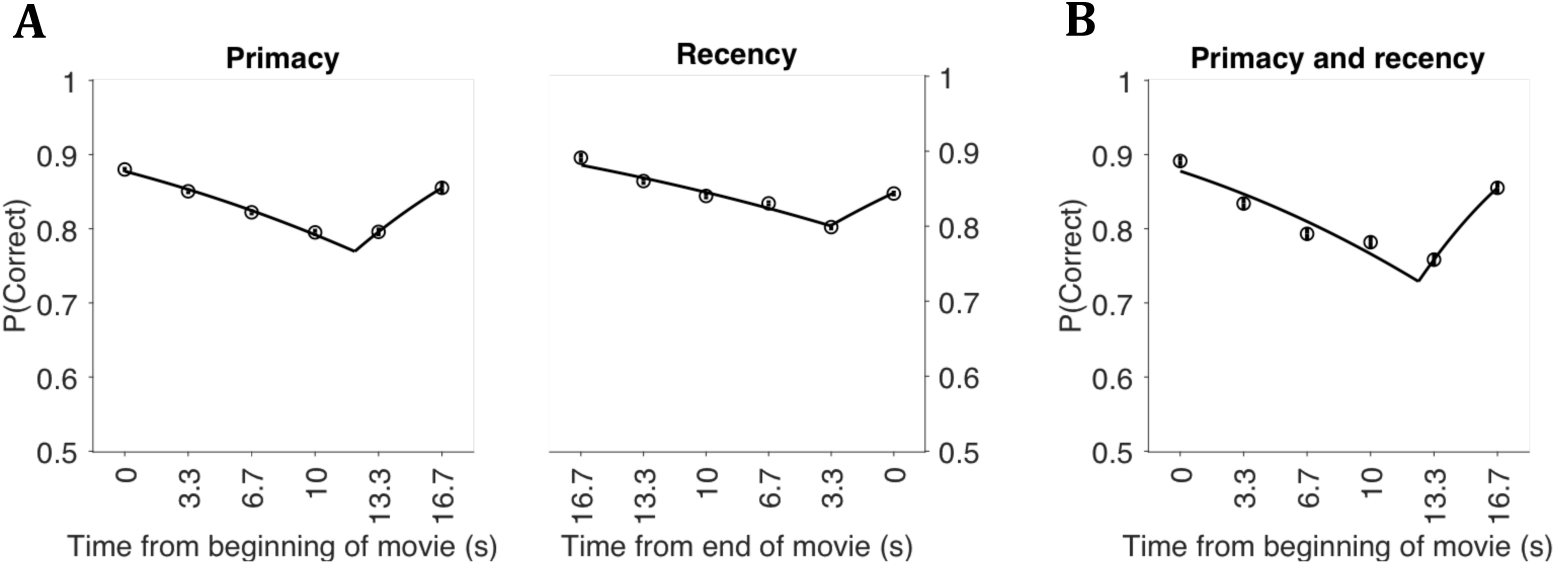
Primacy and recency influenced retrieval accuracy. **(A)** Retrieval accuracy as a function of the time in the movie probed in the retrieval trials. Data points show average accuracy across subjects for times aligned to the beginning (left panel) or end (right panel) of the movie. Lines are model fits of Eq. 5. All retrieval trials from both morph and non-morph blocks were combined across all movie lengths. **(B)** Same as A, but for blocks with 6 feature movies. Because a single movie length is used in this panel, alignment to the beginning or end of the movie are the same. Error bars are SEM.

However, the strength of primacy and recency effects varied considerably for different memory components (Figure 4). Whereas time-mismatch trials were strongly influenced by both effects (primacy: *β*_1_ = 0.22 ± 0.024, p < 10^−8^, recency: *β*_2_ = 0.31 ± 0.07, p = 1.9×10^−5^; FWER-corrected, Eq. 5), primacy and recency were weak or virtually absent in feature-mismatch (primacy: *β*_1_ = - 0.094 ± 0.61, p = 1.0, recency: *β*_2_ = 0.36 ± 0.17, p = 0.064; FWER-corrected Eq. 5), location-mismatch (primacy: *β*_1_ = 0.22 ± 0.34; p = 1.0, recency: *β*_2_ = 0.075 ± 0.036, p = 0.076; FWER-corrected, Eq. 5), and novel-feature trials (primacy: *β*_1_ = 0.17 ± 0.16, p = 0.6, recency: *β*_2_ = 0.026 ± 0.11, p = 1.0; FWER-corrected Eq. 5). Primacy for time-mismatch trials was consistent across individual subjects, with 9/10 showing positive primacy slopes (Fig. S2C; *β*_1_>0; p=0.011, one-way sign test, Eq. 5). Recency for time-mismatch trials was not as consistent with only 7/10 subjects showing positive slopes (Fig. S2D; *β*_2_>0; p=0.17, one-way sign test, Eq. 5).

**Figure 4.**
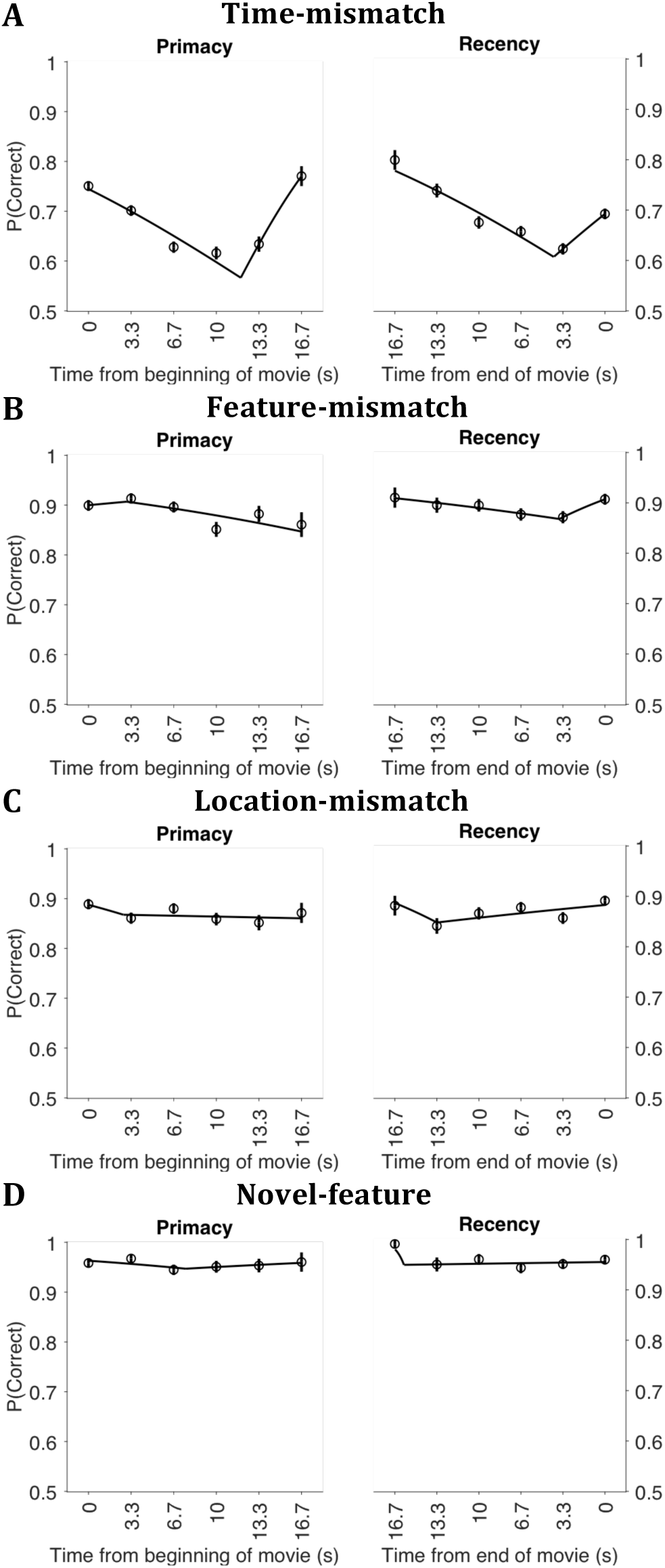
Primacy and recency were most pronounced for time-mismatch trials. Conventions are as in Fig. 3A.

To quantitatively compare the susceptibility of different memory components to primacy and recency, we focused on blocks with the longest movies, where primacy and recency were strongest (Fig. 5). The slope of the reduction of accuracy for intermediate features compared to earlier features was significantly more positive on time-mismatch trials than location-mismatch trials (stronger temporal primacy; *β*_4_ = 0.43 ± 0.12, p = 0.00084; Eq. 6, FWER-corrected). An effect in the same direction existed when comparing time-mismatch to feature-mismatch trials too, although it did not reach statistical significance (*β*_4_ = 0.23 ± 0.15; p = 0.36; Eq. 6, FWER-corrected). Also, the slope for the increase of accuracy for latest features compared to the intermediate features was significantly steeper on time-mismatch trials for both comparisons (stronger temporal recency; compared to location-mismatch trials: *β*_5_ = 0.33 ± 0.12; p = 0.016; compared to feature-mismatch trials: *β*_5_ = 0.34 ± 0.13; p = 0.033; Eq. 6, FWER-corrected). In contrast, the strength of primacy or recency was comparable for location-mismatch and feature-mismatch trials (Fig. 5C; Primacy: *β*_5_= - 0.094 ± 0.092; p = 0.93; Recency: *β*_5_ = 0.17 ± 0.41; p = 1.0; Eq. 6, FWER-corrected). Overall, our results suggest that the “when” component of episodes was most influenced by primacy and recency.

**Figure 5.**
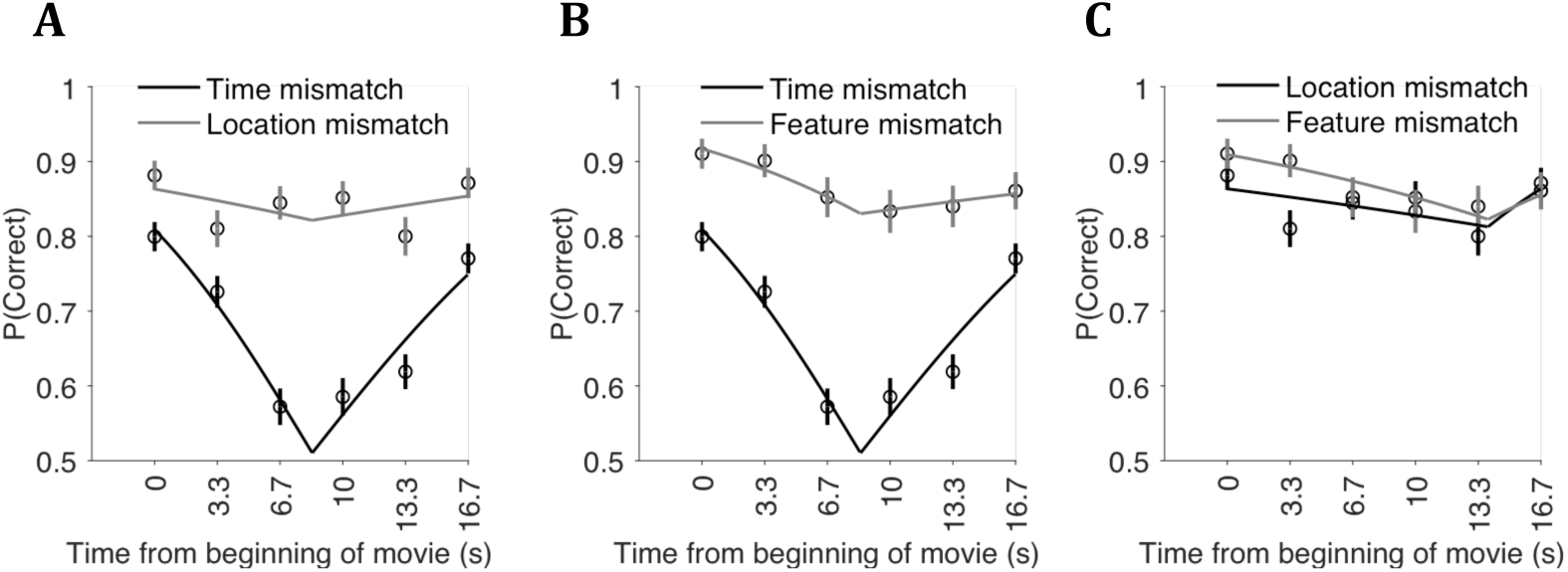
Quantitative comparison of the susceptibility of different memory components to primacy and recency. Data are from blocks with six-feature movies. In each panel, black represents one trial type and gray represents another trial type (see legends). Lines are fits to Eq. 6. Primacy and recency slopes are larger for time-mismatch trials, but comparable for feature- and location-mismatch trials. Plotting conventions are as in Figure 3B.

### Increasing mnemonic load differentially affects the accuracy of different components of episodes

In order to further probe the separability of the WWW components of episodes, we asked if we increased the load on one of the components, would each of them be impacted equally? To test this, we used a variant of our task in which the movie included two features with parametrically variable similarity. One of the features was a mixture of another feature in the movie with a third feature that was not shown in the encoding phase. By changing the mixing coefficient (Eq. 1), we could make the feature pair (the “morph pair”) similar and hard to discriminate (Fig. 6), or distinct and easy to discriminate, or anything in between (Fig. S4A, Methods). Overall, 58% of the blocks in a session showed a morph pair in the encoding phase (see Methods for more details). Because of the presence of other features in the movie, only a fraction of retrieval trials (49%) probed the morph pair in those blocks. However, subjects could not know while watching a movie whether the morph pair would (or wouldn’t) be in the retrieval trials. The best strategy was therefore to discriminate and memorize the morph pair as well as other features in the movie. We hypothesized that blocks with two similar features in the movie imposed a higher mnemonic load, which could interfere with the storage and subsequent recall of *all* features.

**Figure 6.**
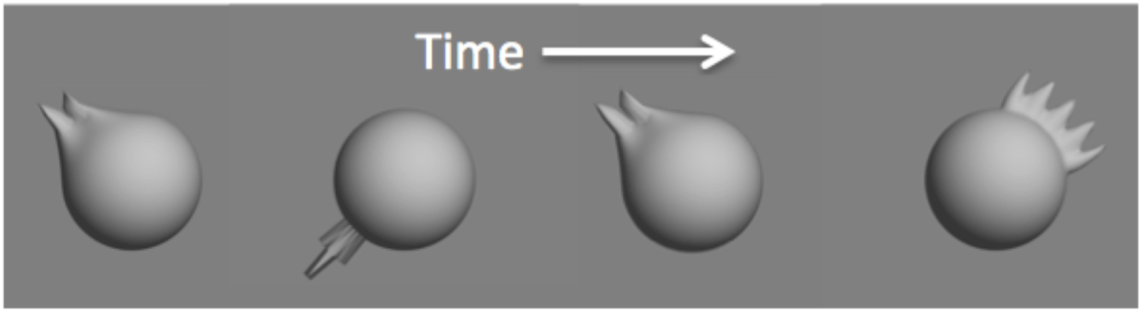
Introduction of morph features. Schematic of example movie cue, as in Fig. 1A inset, but from a morph block. Note the similarity between the 1^st^ (morph level = 0.25) and 3^rd^ (morph level = 0.5) features.

Could the presence of the morph pair in the movie interfere with accuracy on retrieval trials that did not specifically probe the morph pair? And, did the increased mnemonic load compromise accuracy on some trial types more than others? The answer to both questions is yes. Focusing on retrieval trials that targeted features other than those in the morph pair, we observed a reduction of accuracy in time-mismatch compared to location-mismatch trials of non-morph features that significantly depended on the similarity of the morph pair (*β*_3_ = −0.91 ± 0.3, p = 0.0078; Eq. 7, FWER-corrected). Further, we found a trend towards morph-pair-dependent lower accuracy on time-mismatch than feature-mismatch trials (*β*_3_ = −0.82 ± 0.35; p = 0.063; Eq. 7, FWER-corrected). However, there was no evidence of a morph-pair-dependent difference between location-mismatch and feature-mismatch trials (*β*_3_ = −0.098 ± 0.39; p = 1.0; Eq. 7, FWER-corrected). Overall, the “when” component of memory was most affected by the increased mnemonic load caused by the morph pair.

Our hypothesis about the effect of memory load on retrieval accuracy also predicts that longer gaps between the morph pair in movies of the same length would cause lower recall accuracy, as a larger number of intervening features (NIF) makes distinction of the morph pair more difficult (in other words, larger NIF should increase memory interference). Figure 7 shows the retrieval accuracy of different trial types as a function of the NIF. As in the previous analysis, we excluded trials that specifically probed the morph pair and focused on the remaining features of the movie. Since movies showed between 3 to 6 features, the possible NIF ranged between 0 (when the morph pair occurred back-to-back) and 4 (six-feature movies with the morph pair shown at the beginning and end). There was a significant decline in retrieval accuracy with increased NIF for time-mismatch (Fig. 7A; *β*_1_ = −0.25 ± 0.046; p = 8.4×10^−8^; Eq. 8, FWER-corrected) and location-mismatch trials (Fig. 7B; *β*_1_ = −0.19 ± 0.058; p = 0.0025); Eq. 8, FWER-corrected). Feature-mismatch trials did not show a significant change in accuracy (Fig. 7C; *β*_3_ = −0.026 ± 0.077; p = 1.0; Eq. 8, FWER-corrected). To test if changes in retrieval accuracy with the NIF were stronger for certain trial types, we performed pairwise comparisons between the slopes in Fig. 7. The drop of accuracy was significantly steeper for time-mismatch trials than feature-mismatch trials (*β*_3_ = 0.26 ± 0.099, p = 0.028; Eq. 9, FWER-corrected). A similar trend existed for comparison of time-mismatch and location-mismatch trials, but it did not reach statistical significance (*β*_3_ = 0.13 ± 0.081; p = 0.30; Eq. 9, FWER-corrected). There was no difference between the location-mismatch and feature-mismatch trials (*β*_3_ = 0.12 ± 0.11; p = 0.72; Eq. 9, FWER-corrected).

**Figure 7.**
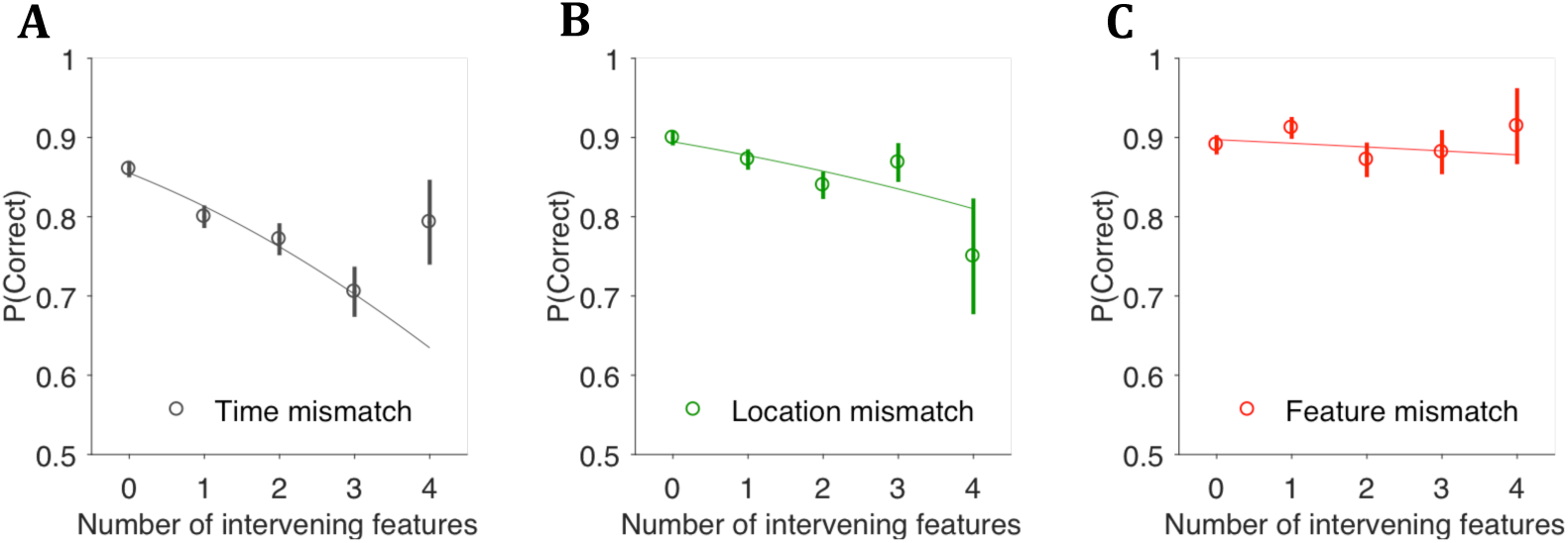
Increasing memory load with larger number of movie features intervening between the morph pair affects the time-mismatch trials most strongly. Time-mismatch (A) and location-mismatch trials (B), but not feature-mismatch trials (C), show a decline in accuracy with increasing the NIF. Lines are fits to Eq. 9. Error bars are SEM.

These results indicate that accuracy on time-mismatch and location-mismatch trials, which test memory for “when” and “where”, was affected by the number of intervening features between the morph pair. Surprisingly, feature-mismatch trials, which test memory for “what,” were minimally affected. The significant difference between these WWW trial types provides additional evidence that encoding and retrieval of different components of episodes are separable.

### Similar performance in a task variation designed to interfere with working memory

The relatively short gap (<1 min) between encoding and retrieval in our main task may raise a concern that subjects performed the task by relying *solely* on working memory. Although recent studies reduce the likelihood of this concern by showing that regions involved in episodic memory also engage in tasks similar to ours^23-25^, we designed a control experiment that directly tested the role of working memory in shaping the results explained in previous sections. This experiment, which disrupted working memory, was a “1-back” version of our TRANSFER task, in which subjects responded to the retrieval trials based on the movie shown in the block before (1-back from) the current block (Fig. S5A; Methods). Therefore, subjects were tasked to hold two movies in mind simultaneously, and responses to retrieval trials were separated from the relevant movie by the retrieval trials from the previous block and the movie of the current block. Five of our ten original subjects performed this task variant.

Despite the high intensity interference of working-memory, subjects had only a small reduction of accuracy for the substantial increase in the complexity of the 1-back task compared to the original TRANSFER task (overall accuracy in non-morph blocks, 83.7±5.5% on the 1-back task vs. 91.0±4.7% for the original task). Critically, the main effects that demonstrated separability of the three memory components on the original task were also present in the 1-back task. Subjects showed unequal accuracies across trial types, which reached statistical significance (p=0.0016, likelihood ratio test, Eq. 2 vs. Eq. 3) despite a much smaller trial count (<20%) in the 1-back task. The average pairwise accuracy differences were the same as the original task (6.2±4.6% vs. 5.1±3.5%). Further, the accuracies of different trial types followed a similar pattern as on the original task (Fig. S5B): time-mismatch trials showed the lowest accuracy (80.5±7.1%), while novel-feature trials showed the highest accuracy (94.6±7.4%). The trial counts in this task variant were too small to test the effect of memory load. However, the primacy (Fig. S5C; *β*_1_ = −0.14 ± 0.086; p = 0.054, Eq. 5) and recency effects (Fig. S5C; *β*_1_ = −0.3 ± 0.15; p = 0.027, Eq. 5) were present and strong. Notably, as with the original TRANSFER task, the strongest serial order effects occurred for time-mismatch trials, which showed significant primacy (Fig. S5D; *β*_1_ = −0.27 ± 0.072; p = 0.0001, Eq. 5) and a trend towards recency (Fig. S5D; *β*_1_ = −0.36 ± 0.25; p = 0.077, Eq. 5).

These similar patterns of accuracies between trial types on the original and 1-back TRANSFER tasks support the hypothesis that the differential mnemonic effects between trial types in the original TRANSFER task were not due to working memory mechanisms. Instead, they are inherent to the encoding and retrieval of short-term episodes.

## Discussion

We developed a task to simultaneously investigate memory for the what, where, and when (WWW) components of episodes. The TRANSFER task did not instruct subjects to focus on any particular WWW component, thereby enabling unbiased comparison of the accuracy of memory components. We found that the WWW components of an episode did not form a coherent whole that was inseparably stored and retrieved. Instead, retrieval accuracies differed across components, with memory for “when” showing the lowest accuracy. Further, the WWW components had different susceptibility to primacy, recency, and interference caused by memory load—three common phenomena associated with memory. Trials probing “when” showed the strongest primacy and recency effects, and trials probing “when” and “where” were most interfered by increasing memory load through a task manipulation that required discrimination of similar features. These results suggest that memory processes maintain (at least) partial separation of the associated WWW components, with the subjective coherence of episodes being a result of active reconstruction from the associated components.

The abstract visual features in the TRANSFER task and instructions that gave subjects no indication of task goals were integral characteristics of our design. Previous work has had difficulty isolating memory for item and order information^18,26,27^, largely owing to the use of sequences of consonants or words that introduced phonemic coding confounds^12^. Later studies used abstract shapes to avoid such confounds. However, their reliance on task instructions that focused subjects’ attention on one or the other memory component^13-17,19^ made it difficult to parse if differences in memory effects were specific to the instructions or inherent to the organization of memory. We overcame this challenge by using abstract shapes and only instructing subjects to identify trials as “match” or “mismatch” during the retrieval phase without directing attention to any particular component.

On first glance, it might seem that the lower accuracy on time-than location- and feature-mismatch trials is merely due to greater difficulty of perceiving “when” in the TRANSFER task. But perceptual difficulty does not explain our results. As mentioned in the introduction, if we took any of the WWW components and separately designed a task to probe only one of them (e.g., to probe when, show two features serially and ask which came first), accuracy would be at or near 100%^20^. In other words, none of the WWW components were near psychophysical threshold for discriminating the “what” of the feature (excluding trials in our morph blocks, Figs. 6-7), “where” it was on the sphere, or “when” it was shown in the movie. Therefore, unequal accuracies should not come from differences in sensory discrimination during the movie. As for the number of each WWW component, 470 features were shown in 3-6 time slots in 4 locations. But in any given movie, the same number of each WWW component had to be remembered (e.g. a 5 feature movie had 5 features in 5 locations in 5 time slots). If anything, since there were only 4 possible locations, location-mismatch trials might be expected to show the lowest accuracy due to intrusions from the frequent reuse of locations both between and within trials^20^. However, despite this potential for redundancy confusion, responses on time-mismatch trials were significantly less accurate than location-mismatch trials (Fig. 2). Therefore, difficulty of time-mismatch trials was due to the inherent challenge in maintaining temporal order. This conclusion is also supported by stronger primacy and recency effects for the time-mismatch trials vs. other trial types (Figs. 4-5). These results suggest that how information about when is encoded and retrieved as part of an episode differs from those that maintain what and where.

Serial order effects have frequently been fit with a perturbation model^28^ in which noise causes confusion between an encoded event and its adjoining events with a probability, *θ*. For example, the third feature of a movie in our task could be remembered either as the second feature or as the fourth feature with a probability of *θ*/2, with multiple perturbations possible before recall. Values of *θ* near 0.05 have typically been found using letter or word recall tasks^29^. The strength of primacy and recency in our time-mismatch trials is compatible with *θ* values of 0.094±0.024 across subjects (mean ± SE), suggesting that serial order effects in our task are more potent than in previous tasks^28,29^. This might be expected, considering that phonemic coding strategies^12^ could be less easily utilized with our abstract features.

An advantage of the TRANSFER task is that it equalizes rehearsal of different memory components, which has been hypothesized as a key factor for the strength of primacy. That is, if subjects rehearse all the observed movie features following each new feature in the encoding phase of our task, earlier features will be rehearsed more than later ones, thereby improving their retrieval accuracies^30,31^. However, such automatic rehearsals, if present, would equally apply to the what, where, and when aspects of each feature, and therefore are unlikely to underlie different strengths of primacy for different memory components in our task (Figs. 4-5). Even if subjects employed a phonemic coding strategy like attempting to name the abstract features and rehearse the names in order, they are equally rehearsing both what and when, and yet we find these components to have significantly different accuracies (Fig. 2) and primacy (Figs 4-5). Future work could attempt to remove the stronger primacy effect for the when component by eliminating the rehearsal period or actively preventing rehearsal^30^ to more accurately quantify the effect of automatic rehearsal in our task.

Our finding that time-mismatch and location-mismatch trials were most affected by the presence of two similar features (the morph pair) in the movie (Fig. 7) could reflect a fundamental limitation of memory. Since we manipulated the feature similarity, which bears on the “what” component of episodes, why do we see stronger effects on the when and where components? Considering the current evidence, we suggest that the stronger effect of the morph pair on when and where is related to separability of these components. In blocks without the morph pair, time-mismatch trials were significantly less accurate than location-mismatch trials, which themselves were less accurate than feature-mismatch trials (Fig. 2). If the memory system has a limited mnemonic reservoir^24,32,33^, the added mnemonic load caused by the morph pair would impact the hardest trials more strongly.

Because of the novelty of the TRANSFER task, the full set of brain structures that underlie the subjects’ performance is yet to be determined. Although we are careful to distinguish the relatively short duration of our task blocks from long-term episodic memories, we hypothesize that our task engages the same structures that underlie longer-term episodes. Two observations support this hypothesis. First, we replicated our findings in a 1-back variant of the TRANSFER task, where the retrieval trials in a block were about the movie observed in the preceding block. Not only were the retrieval trials separated from their corresponding movie by a much longer interval in the 1-back task, but also this longer interval included multiple events—the movie of the current block and the retrieval trials of the preceding block—that should strongly interfere with working memory and thwart the ability to rehearse movies before the retrieval trials. Replication of our main findings in the 1-back task rules out mechanistic explanations based solely on working memory, necessitating the involvement of longer-term memory mechanisms^24^. Second, past studies using tasks with short timescales but large memory loads that seem to overwhelm working memory appear to recruit medial temporal lobe structures^23,24,34^. Both the main TRANSFER task and its 1-back variant fall in this category.

“I meet you. I remember you…Why not you in this city and in this night, so like other cities and other nights you can hardly tell the difference?” Elle’s soliloquy in Marguerite Duras’s *Hiroshima Mon Amour* laments on the fungibility of our episodic memories, where it is often difficult for us to separate events in time when they share similar components. In our work, when humans are tasked to remember what, where, and when components of short-term episodes, we find comparable difficulties. In particular, we find temporal information to be the least accurate and most susceptible to interference by serial order and memory load, especially when the recalled episode is surrounded by a sequence of similar episodes. These differences in accuracy between the what, where, and when components of episodes in our experiments suggest a degree of independence between memory components, which speaks against holistic mechanisms that inseparably bind episodes. They also suggest that the apparent coherence of recalled events is largely a product of a reconstruction process^35^ that re-joins the memory components. Future work following up on our results will offer a window into the mechanisms behind the separability of the memory components and how the brain reconstructs the seamless episodes we experience on an everyday basis.

## Methods

Ten human subjects (5 male, 5 female; age, 23.4±4.6 years, mean±s.d.) with normal or corrected-to-normal vision participated in the experiment. All subjects were naïve to the purpose of the experiment, provided written consent before participation, and were free to leave the study at any time. They received fixed monetary compensation for their participation. All procedures were approved by the Institutional Review Board at New York University.

### Behavioral task

During the experiment subjects were seated in an adjustable chair in a dimly lit room with chin and forehead supported 57 cm in front of a Cathode Ray Tube display monitor (20”, EIZO FlexScan T966; refresh rate 75 Hz, screen resolution 1600 × 1200). Stimuli were presented using Psychophysics Toolbox^36^ and Matlab (The MathWorks, Inc.). Eye movements were monitored at 1 kHz with an infrared camera (Eyelink, SR-Research).

An outline of the TRANSFER task is shown in Figure 1A. Each block of trials started with the subject looking at a fixation point (FP, 0.15° radius) at the center of the screen. After a short delay the encoding phase of the block began with the presentation of a movie centered on the screen (Figure 1B). The movie started with a 1.3° radius gray sphere on which 3-6 shapes (“features”) grew and then receded from 4 potential locations on the surface of the sphere (45°, 135°, 225°, or 315° along the circumference). Subjects were required to maintain fixation within 1.5° from the FP throughout the movie. Each feature extended up to 2° from the edge of the sphere in 0.333 s, remained extended for 0.667 s, and then retracted in 0.333 s (total feature duration, 1.333 s). A 2 s delay showing only the sphere was presented after each feature during which subjects maintained fixation but were permitted to blink to prevent dry eyes. Features from each movie were chosen randomly from a pool of 470 unique features. This large pool ensured that most features were not repeated in a day (∼400 features per day), minimizing interactions between movies from different blocks of trials. Subjects were instructed to utilize the 2 s interval after each feature to memorize and rehearse it. In early data collection blocks, an orange progress bar was shown 3° below the FP. The progress bar grew horizontally at a constant speed throughout the movie (0.3°/s) and indicated the time of each feature. In later blocks of data collection, the progress bar was eliminated for all subjects and they were asked to mentally keep track of elapsed time. Removing the progress bar ensured that subjects could not perform the task by visually associating the length of the bar with simultaneously observed features. After the movie, subjects moved from the encoding phase to the retrieval phase, where they were tested on their memory of the movie.

Subjects initiated each retrieval trial by fixating on the FP. After a short delay (0.2-0.5 s, truncated exponential, mean=0.3 s), a blue and a red target (equiluminant, 18.5 cd/m^2^) appeared to the right and left of the FP (0.5° radius, 8° eccentricity). The location of the two targets swapped randomly across trials. After a second short delay (0.2-0.5 s.; truncated exponential with mean=0.3s), a “time cue” appeared on the screen indicating that subjects would be queried about a particular time in the movie. Subjects were instructed to recall the feature shown during the cued time. After a variable delay (0.5-1.5 s.; truncated exponential with mean=0.83 s), an image was shown for 0.3-1.2 s (truncated exponential with mean=0.6 s) at the center of the screen. Subjects decided whether this image matched the cued time in the movie. Finally, the FP, stimulus, and time cue disappeared, indicating that subjects should report their choice with a saccadic eye movement to one of the two targets.

Subjects were instructed to identify “match” or “mismatch” trials during the retrieval phase based on the content they saw during the encoding phase. Choosing the blue target indicated that the image matched the cued time in the movie, and the red target indicated a mismatch. Subjects received distinct auditory feedbacks for correct and error responses. We limited the number of retrieval trials per movie block to minimize potential interference due to consecutive retrievals. For movies with 3, 4, and ≥5 features, we showed 1, 1-2, and 2-3 retrieval trials, respectively. To avoid learning from feedback of previous retrieval trials in a block, each cued time in the movie could be queried only once in the retrieval phase.

For a retrieval image to be a match, it should include the same feature at the same location as those shown in the movie at the cued time. A mismatch could arise for different reasons. Throughout the paper we focus on four mismatch trial types: “**location-mismatch**,” where the retrieval image was a movie frame of a time-matching feature shown in a different location than during the movie; “**feature-mismatch**,” where the retrieval image was a movie frame of a mismatching feature taken from a different time in the movie but shown in a time-matching location; “**time-mismatch**,” where the retrieval image was a movie frame that appeared at a time other than the cued one; and “**novel-feature**,” where the retrieval image consisted of a feature not shown during the movie (but appearing at a time-matching location). The match and mismatch trials were balanced, each comprising 50% of retrieval trials. Location, feature, and time-mismatch trials were balanced too, each comprising 30% of mismatch trials (15% of all trials). Novel-feature trials comprised the remaining 10% of mismatch trials (5% of all trials). The match and mismatch trials were randomly intermixed throughout the session.

By the nature of the task, time-mismatch trials always showed the retrieval image of a feature in the original location it was shown in the movie, but with a cue indicating a different time slot from the movie. However, for feature-mismatch trials, since the retrieval image was selected as one of the other features from a different part of the movie, there was a ¼ chance (due to there being only 4 possible locations) that the location of the other feature would happen to be the same location as the original feature shown during the cued time slot. In that case, since the subject would see a feature from a different time of the movie, but in the same location it was originally shown, the retrieval image was functionally identical to a time-mismatch trial. Therefore, in our analyses we grouped these trials with our time-mismatch trials.

To ensure that consecutive movie blocks did not lead to confusion and interference, the retrieval trials of a block were separated from the movie of the next block with a 3 s interval during which the screen turned blue to demarcate the end of the previous block. Subjects were instructed to rest and prepare for the next block.

Subjects were trained until they achieved a 75% accuracy criterion (typically for 3-4 days). Data collection began following training. Subjects typically performed the task for one hour each day, during which they completed 6-8 sessions (5-8 min per session), each including 15 blocks. Each subject contributed 2390-4877 trials to the dataset for the TRANSFER task (median, 3624 trials; 14-28 hours of data collection per subject). Additionally, 5 of the 10 subjects contributed 1216-2097 trials for the 1-back variant of the TRANSFER task explained below (median, 1466 trials; 10-14 hours of data collection per subject). Combined, there were 38,735 trials for the TRANSFER task, and a separate 7,416 trials for the 1-back TRANSFER task.

### Stimuli

All features were parametrically defined as protrusions of a fixed number of vertices on the surface of a sphere. We started with 62 prototype features that were manually created to be easily discriminable. Next, to increase the number of features such that subjects could not easily memorize the entire set, we took each possible pair of these 62 prototype features and combined them such that the vector of vertices for each feature was exactly halfway between the pair (as explained in the next paragraph). This created 1922 unique features. We used the 470 of these that displayed a smooth transition when one prototype was morphed into the other. We call these 470 our main feature set.

Parametric morphing of features was implemented according to the following equation:

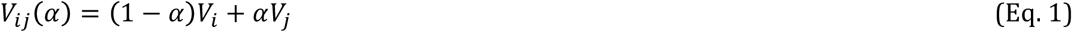

where *V*_*i*_ and *V*_*j*_ are the vectors of vertices for any of our 470 pairs of prototypes *i* and *j, α* is the mixing coefficient that defines the morph level, and *V*_*ij*_ (*α*) is the vector of vertices of the intermediate morph between the two prototypes. For the main feature set, the morph level *α* was set to 0.5.

However, by changing the morph level between 0 and 1, we could also parametrically morph one prototype into another (Fig. S5). As we lower the value of the morph level from 0.5 towards 0 (or, symmetrically, raise it from 0.5 to 1), we create a morphed feature with increasing dissimilarity from one of the 470 main features (and increasing similarity to one of the 62 prototype features, Fig. S5). We used such parametrically morphed features in a subset of task blocks as explained in the next section.

### Task variations

#### Parametric variation of feature similarity

To test how the similarity of features in the movies influenced behavior, we included one parametrically morphed feature in the movie in some blocks (67% of blocks for 6 subjects and 50% of blocks for the other 4 subjects). The feature was made by selecting one of the 470 main features and decreasing the value of the morph level from 0.5, as explained in the previous section (Eq. 1). Both the parametrically morphed feature and its partner in the main set, which we refer to as the morph pair, were included in the movie. Therefore, for mixing coefficients close to 0.5, the movie contained two similar features, whereas for mixing coefficients close to 0, the morph pair were visually distinct and easily discriminable (Fig. S4). The value of the mixing coefficient in different blocks was chosen randomly from a truncated exponential distribution with the highest probability for *α* = 0.5 − *ϵ*, and the lowest probability for *α* = 0 (mean *α*, 0.33). *α* = 0.5 was excluded to ensure an objectively correct answer on every trial. Because of this exponential distribution, many of the morph pairs were hard to distinguish (Fig. S4). The order and time of the two features of the pair varied randomly in the movies across blocks. However, the locations of the pair were always the same in a movie. This location varied in different blocks.

#### 1-back TRANSFER task

To test whether subjects’ performance depended on the immediate working memory, we created a variant of the TRANSFER task in which retrieval trials in a block did not probe subjects’ memory of the movie in the same block. Rather, they referred to the movie observed in the preceding block (1-back). The session began with subjects viewing a movie, then a second movie, followed by the retrieval trials referencing the first movie, then a third movie followed by the retrieval trials referencing the second movie, and so on (Fig. S5). To perform this task, subjects had to commit the movie of the current block to memory and then retrieve the memory of the movie from the previous block, overcoming two sources of interference: the immediately observed movie and the retrieval trials that followed the relevant movie. Task parameters remained the same as the main task, with the exception that we limited the length of the movies to 3-4 features. To remind subjects that the retrieval trials are not related to the immediately observed movie in the current block, the color of the time cue of the retrieval trials was distinct from the color of the progress bar of the movie of the current block, but it matched that of the movie of the previous block. 5/10 subjects performed this task variation.

### Data analysis

We tested our hypotheses using a series of regression analyses. Regression coefficients were estimated using maximum likelihood methods. Standard errors of coefficients were estimated by inverting the Hessian matrix of the model log-likelihood function and calculating the square root of the diagonal elements of the inverted matrix^37^. For maximum likelihood fitting, we derived the closed form of the Jacobian and Hessian matrices wherever possible. Where we could not directly derive the matrices, we estimated them numerically. Because single subject’s data were consistent with each other (see Figs. S1-S2), we pooled trials across subjects for all statistical tests. We also report single subject results to provide a complete picture of the strength and diversity of results.

To compare accuracy between time-mismatch and other trial types we used the following logistic regression:

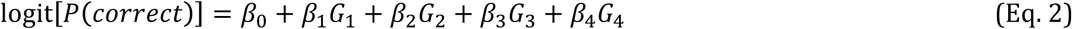

where *β*_*i*_ are regression coefficients and *G*_*i*_ are indicator variables for different trial types. *G*_1_ is for match trials, *G*_2_ is for location-mismatch trials, *G*_3_ is for feature-mismatch trials and *G*_4_ is for novel-feature trials. Eq. 2 served two purposes. First, it enabled us to perform pairwise comparisons between the accuracy of each of the four trial types indicated by *G*_1−4_ and the time-mismatch trials, whose accuracy is captured by *β*_0_. Second, this equation was used to test whether the null hypothesis of identical accuracy for all five trial types adequately explains the data. For this purpose we use a likelihood ratio test^38^ against a logistic regression that captures the null hypothesis that subjects’ accuracy was the same for all trial types:

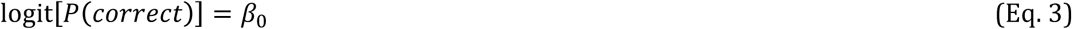

We also reported a modified version of Eq. 2 without the *β*_4_ factor to ensure the differences in accuracies were not driven solely by the novel-feature trials but were present for the three WWW components. For the pairwise comparisons between trial types, we corrected for family-wise error rate using the Bonferroni correction. Therefore, where indicated in the results, p-values were Bonferroni-corrected using the number of comparisons done for that test and significance was considered to be p<0.05.

To compare accuracy between novel-feature and other trials types, we used a modified version of Eq. 2:

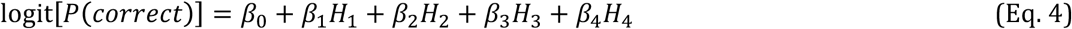

where *β*_0_ represents accuracy for novel-feature trials and *H*_1_ is for match trials, *H*_2_ is for time-mismatch trials, *H*_3_ is for location-mismatch trials and *H*_4_ is for feature-mismatch trials.

Subjects’ performance in the retrieval trials was influenced by both primacy and recency. Primacy caused features closer to the beginning of a movie to be remembered more accurately. Recency caused features closer to the end of the movie to be recalled more accurately. The combination of the two effects led to a v-shaped deflection in the plots that show retrieval accuracy as a function of cued time in the movie (Figs. 2-4). We used the following logistic regression to quantify the strength of primacy and recency:

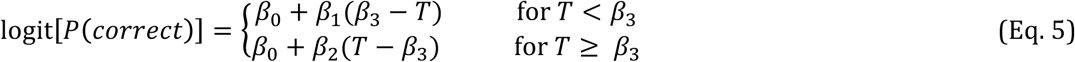

where *T* is the cued time in the movie and *β*_*i*_ are regression coefficients. *β*_3_ defines the time in the movie when a phase transition from a primacy-dominant regime (first line of Eq. 5) to a recency dominant regime (second line of Eq. 5) occurred. *β*_1_ and *β*_2_ quantify how quickly accuracy changed as a function of the cued time in the movie (the “slope”). We use these slopes to report the strength of primacy and recency, respectively, with significant positive slopes indicative of systematically higher accuracy for features shown at the beginning or end of the movie, respectively. To avoid local maxima, we repeated each fit 500 times from random starting points and chose the fit with the highest log likelihood.

Because of the variable movie lengths across blocks, we align feature presentation times to either the beginning or end of the movie and fit Eq. 5 separately for each alignment. The reported primacy effects are based on *β*_1_ of data aligned to the beginning of the movie and the reported recency effects are based on *β*_2_ of data aligned to the end of the movie. This was not necessary for Fig. 3B, since when only one movie length was used, primacy (*β*_1_) and recency (*β*_2_) slope can be measured from the same fit. When fitting Eq. 5 to individual trial types, we once again corrected for family-wise error rate for both primacy and recency slopes by Bonferroni-correcting the p-values using the number of comparisons (N=4) done for each. We also report a one-way sign test across individual subjects, where we assess significant primacy or recency by fitting *β*_*i*_ for each subject’s data with the null hypothesis that the subjects’ slopes will not be different than a distribution with a median of 0.

The discontinuity in Eq. 5 prohibited calculation of the standard errors when *β*_3_ approached an integer multiple of the inter-feature interval in the movies. To obtain an approximate standard error in such cases, we jittered the estimated value of *β*_3_ by 2% to move the log-likelihood function to a continuous region where the calculation of standard errors was permissible (i.e., diagonals of the inverse Hessian were positive). In all cases we confirmed that the small jitter of *β*_3_ minimally changed the overall log-likelihood of the model (maximum change <1%). We also confirmed that the jitter minimally influenced the other regression coefficients. To quantify the effects of jitter on other parameters, we kept *β*_3_ fixed at the jittered value and fit the other coefficients. The jitter led to <1% change in other coefficients.

To compare the magnitude of primacy and recency between two different trial types, we used a modified version of Eq. 5:

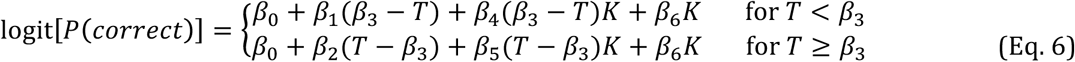

where *K* is an indicator variable for trial type (0 for one type and 1 for another). *β*_4_ and *β*_5_ define the difference in primacy and recency slopes between the two trial types, respectively. *β*_6_ accounts for the overall difference in accuracy between the two trial types. The null hypothesis was no difference in primacy or recency. Since we did 3 tests, p-values were Bonferroni-corrected for family-wise error rate.

Movies in morph blocks contained a morph pair, which was intended to increase the mnemonic load on “what” memory by tasking subjects to differentiate two similar features (see Task Variations). We quantified the effect of morph feature similarity on the accuracy between different trial types using the following logistic regression:

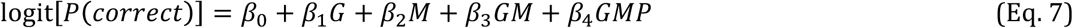

where *G* is an indicator variable that contrasts two retrieval trial types—for example location-mismatch vs. feature-mismatch trials, *M* is the morph level, ranging from 0.5 to 0 on morph blocks and 0 on non-morph blocks, and *P* is an indicator variable that is 1 when the cued time of the retrieval trial probes one of the morph pair in the movie and 0 otherwise. *β*_0−4_ are regression coefficients, where *β*_0_ accounts for the accuracy of one trial type and *β*_1_ accounts for the difference of accuracy with the other trial type, *β*_2_ accounts for the overall effect of the similarity of morph pair on the accuracy of the two trial types, and *β*_3_ quantifies how much the difference in accuracy between the two retrieval trial types is influenced by the similarity of the morph pair. *β*_4_ controls for trials that involve the morph pair, ensuring that *β*_3_ isolates the effect for the retrieval trials that do not involve the morph pair. The null hypothesis of interest is no difference of accuracy between trial types due to the morph pair (*H*_0_: *β*_3_ = 0). Results were FWER-corrected for the three tests used.

In morph blocks, the morph pair could be shown back to back in the movie or be separated by one or more distinct features. The temporal separation of the morph pair in the movie could change memory load via both elapsed time and interference from intervening features. To test whether the number of intervening features (NIF) between the morph pair influenced subjects’ accuracy, we used the following logistic regression:

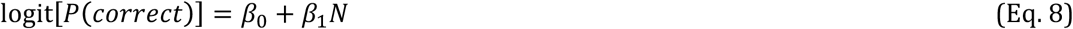

where N is the number of intervening features. The null hypothesis was that the number of intervening features did not influence accuracy (*H*_0_: *β*_1_ = 0).

To test whether the NIF had a differential effect on the memory of location, feature, or time, we extended Eq. 8 to:

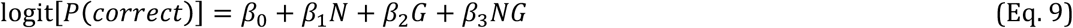

where *G* is the indicator variable for a pair of trial types. *β*_3_ quantifies whether the accuracy difference of the pair of trial types is modulated by the NIF. The null hypothesis was no modulation (*H*_0_: *β*_3_ = 0).

## Acknowledgements

We thank Gouki Okazawa, Michael Waskom, Saleh Esteki, and Mike Kahana for helpful discussions. This research was supported by the National Institutes of Health (R01-MH109180), Simons Collaboration on the Global Brain (542997), and a Pew Scholarship in the Biomedical Sciences.

## Author Contributions

JJS and RK designed the experiment and analyses. JJS collected the data and performed the analyses. JJS and RK wrote the paper.

## Data availability

Full data is available upon request.

## Code availability

Analysis code (in Matlab) and pre-processed data to reproduce figures is available upon request.

---

**Figure S1.**
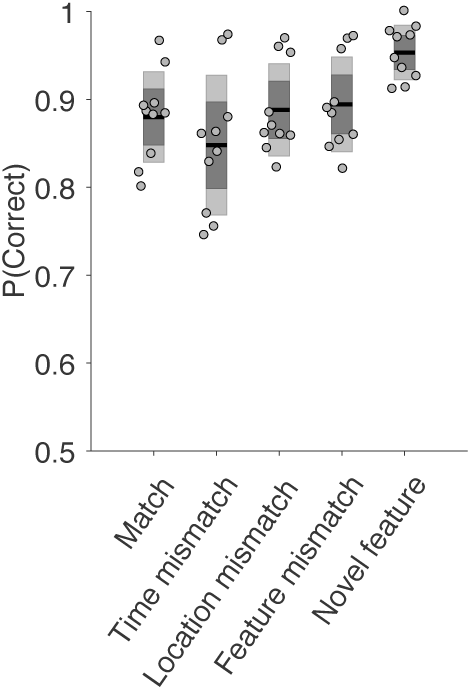
Retrieval accuracy for individual subjects split by trial types. Dots show individual subject’s accuracies and are jittered horizontally for clarity. Black lines are the average for each trial type, dark gray bars are 95% confidence intervals, and light gray bars are standard deviations.

**Figure S2.**
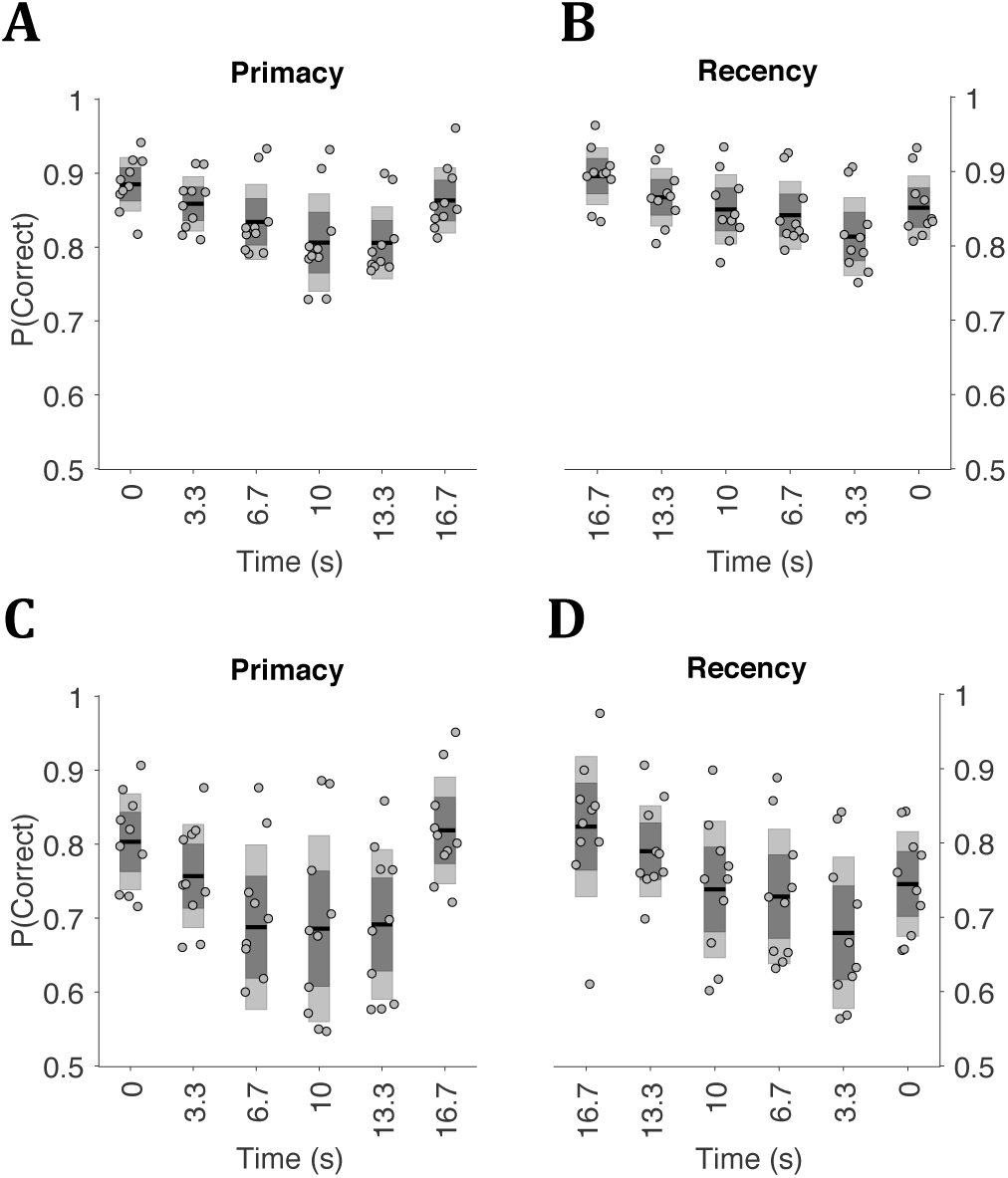
Primacy and recency plots for individual subjects. Retrieval accuracies are plotted as a function of the time of the cued feature in the movie. Dots show individual subject’s accuracies and are jittered horizontally for clarity. Black lines are the average for each cued time, dark gray bars are 95% confidence intervals, and light gray bars are standard deviations. Data are combined across blocks with different movie lengths. **A)** Aligned to beginning of the movie, combined across all trial types. **B)** Aligned to end of the movie, combined across all trial types. **C-D)** Same as **A-B**, but for time-mismatch trials only.

**Figure S3.**
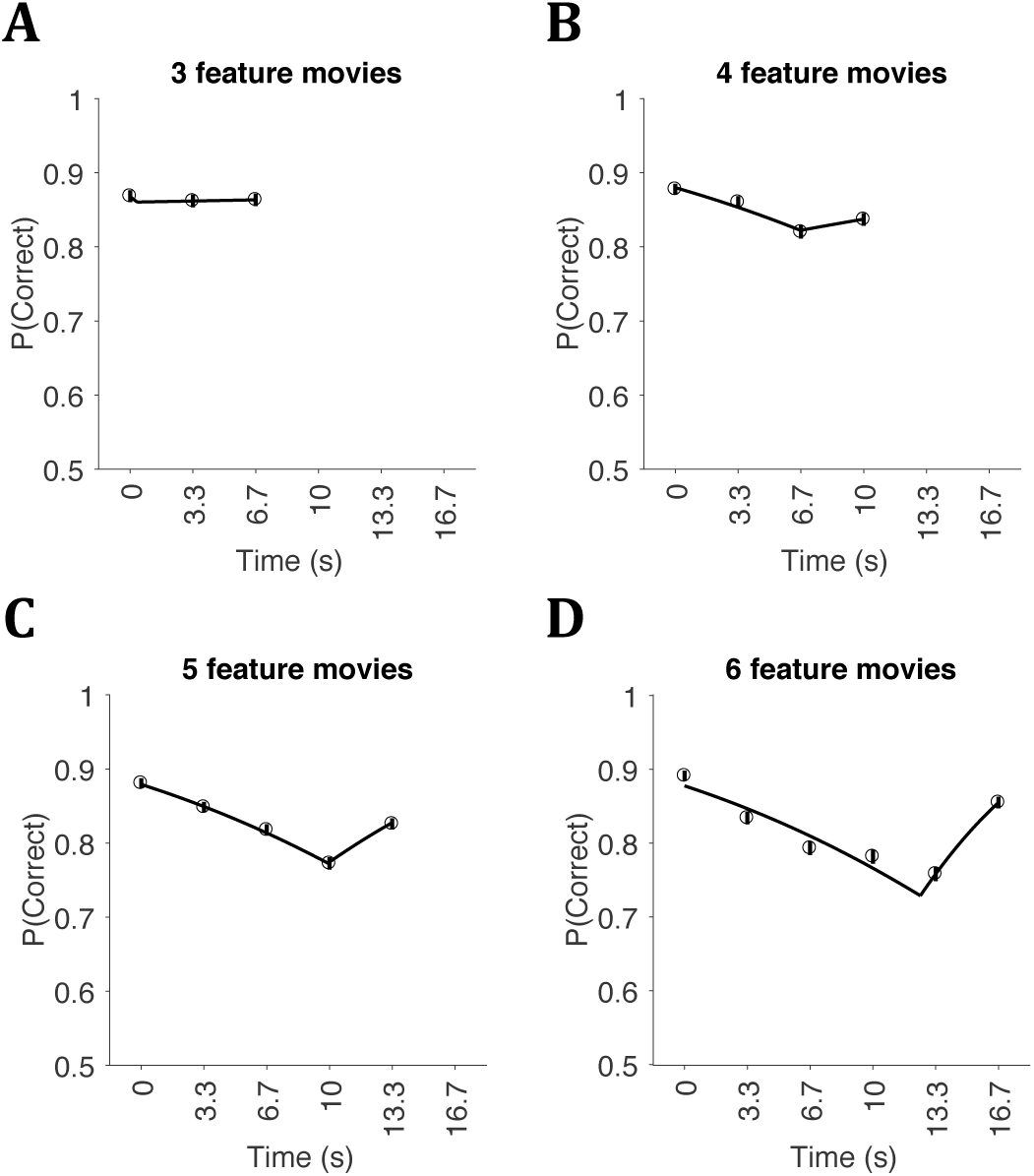
Primacy and recency effects strengthen in trials from longer movies. All conventions are similar to Figure 2B. **A-D** show trials grouped from movies of particular lengths, from 3 to 6 features, respectively. Primacy slope is significantly different from 0 for movie lengths of 4 or more (Eq. 5, movie length 3: *β*_1_ = 0.072 ± 0.21; p = 0.36; movie length 4: *β*_1_ = 0.21 ± 0.084; p = 0.0065; movie length 5: *β*_1_ = 0.26 ± 0.043; p < 10^−8^; movie length 6: *β*_1_ = 0.28 ± 0.029; p < 10^−8^). Recency slope is significantly different from 0 for movie lengths of 5 and 6 (Eq.5, movie length 3: *β*_2_ = 0.58 ± 2.3; p = 0.4; movie length 4: *β*_2_ = 0.12 ± 0.098; p = 0.11; movie length 5: *β*_2_ = 0.29 ± 0.077; p = 7.6×10^−5^; movie length 6: *β*_2_ = 0.62 ± 0.087; p < 10^−8^).

**Figure S4.**
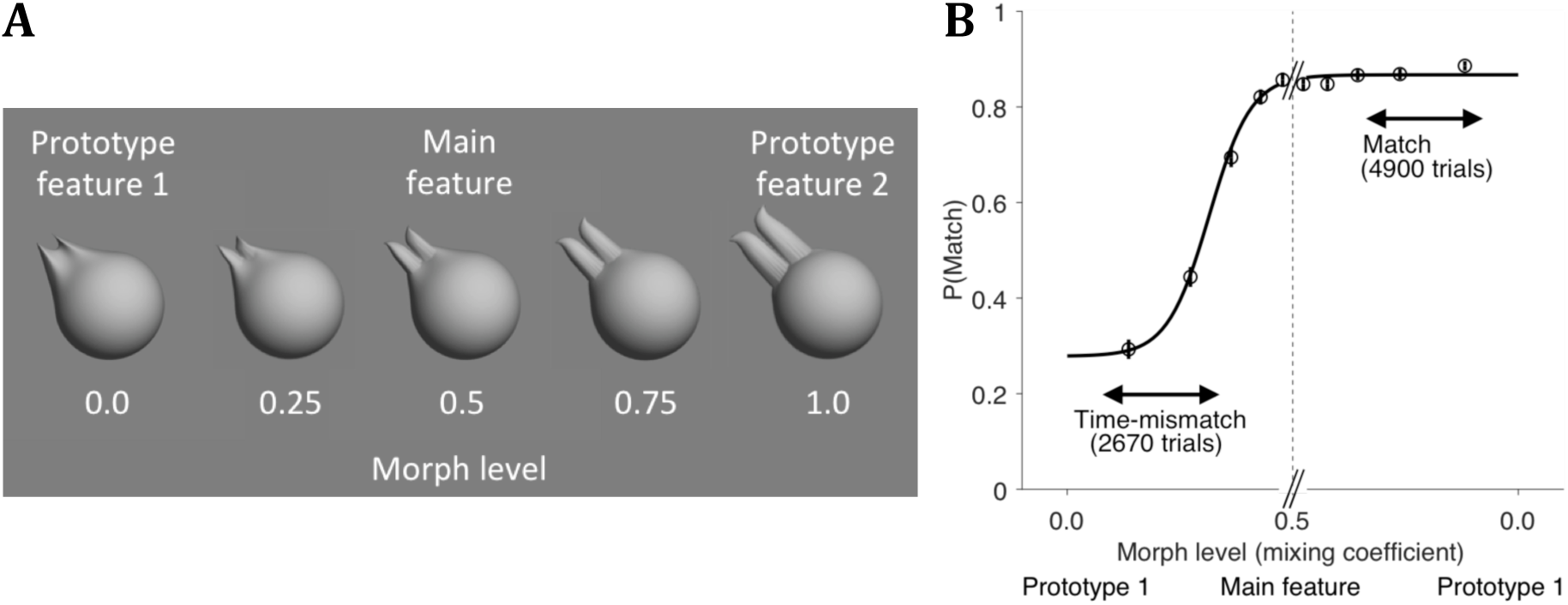
Morphing method and subjects’ accuracy on morphed features. **A)** An example morph line between two of the 62 prototype features. The middle feature with 0.5 mixing coefficient is one of the 470 main features used in the task. A morph pair in the morph blocks comprised of one main feature and a morph between that feature and one of the two prototype features that made it. For example, morph levels of 0.25 and 0.5 correspond to the morph pair shown in Figure 6. **B)** Psychometric plot of the proportion of “match” responses on match and time-mismatch trials in which both the cued time and the retrieval image were associated with the morph pair. For match trials, this meant that the cued time and retrieval image both probed one of the morph pair. For time-mismatch trials, this meant that the time cue referred to one of the morph pair, while the retrieval image showed the other. The five data points on each half of the plot (time-mismatch and match) divide the trials into bins with equal trial counts based on the mixing coefficient of the morph pair. For the time-mismatch trials, the probability of match choices declined as the morph pair became more distinct (i.e., subjects were more likely to distinguish the pair). For the match trials, the probability of match choices increased only slightly with distinctness of the morph pair, probably due to a ceiling effect on the probability of correct responses caused by the overall task difficulty. All morph blocks of the main task are included in this plot. The black solid line is a logistic regression fit.

**Figure S5.**
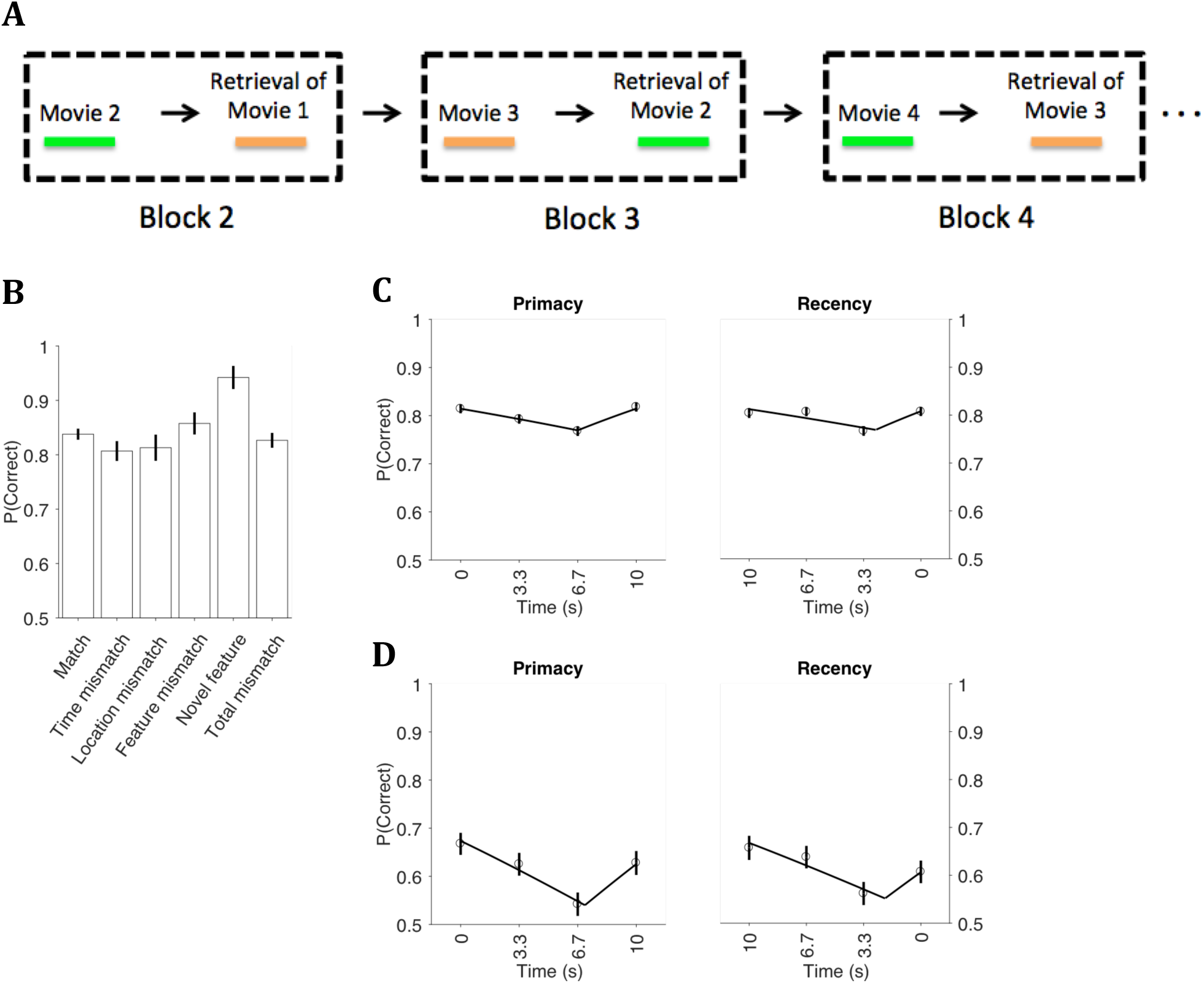
1-back TRANSFER task shows similar accuracies across trial types and serial order effects as the original TRANSFER task. **A)** Design of the 1-back version of TRANSFER task. In the encoding phase of each block, subjects viewed a movie, similar to the original task, but in the retrieval phase, they were asked to answer based on the movie viewed in the preceding block. Movies were limited to 3 or 4 features in length. The progress bar in the encoding phase of odd and even blocks had distinct colors (orange on odd blocks and green on even blocks). The time cues in the retrieval trials matched the color of the progress bar of the corresponding movie to help subjects not confuse the relevant movie. **B)** Accuracy for different trial types of the non-morph blocks in the 1-back task. Conventions are the same as Fig. 2. **C-D)** Primacy and recency for the 1-back task. Conventions are the same as Fig. 3A. C shows trials pooled across all trial types and movie lengths. D shows time-mismatch trials only.

## References

1 Moscovitch, M., Cabeza, R., Winocur, G. & Nadel, L. Episodic Memory and Beyond: The Hippocampus and Neocortex in Transformation. Annual Review of Psychology, Vol 67 67, 105-+, doi:10.1146/annurev-psych-113011-143733 (2016).

2 Nyberg, L. et al. General and specific brain regions involved in encoding and retrieval of events: What, where, and when. P Natl Acad Sci USA 93, 11280–11285, doi:Doi 10.1073/Pnas.93.20.11280 (1996).

3 Tulving, E. Episodic memory: From mind to brain. Annu Rev Psychol 53, 1–25, doi:DOI 10.1146/annurev.psych.53.100901.135114 (2002).

4 Allen, T. A. & Fortin, N. J. The evolution of episodic memory. P Natl Acad Sci USA 110, 10379–10386, doi:DOI 10.1073/pnas.1301199110 (2013).

5 Babb, S. J. & Crystal, J. D. Episodic-like memory in the rat. Curr Biol 16, 1317–1321, doi:10.1016/j.cub.2006.05.025 (2006).

6 Clayton, N. S. & Dickinson, A. Episodic-like memory during cache recovery by scrub jays. Nature 395, 272–274, doi:Doi 10.1038/26216 (1998).

7 Eichenbaum, H., Fortin, N. J., Ergorul, C., Wright, S. P. & Agster, K. L. Episodic recollection in animals: “If it walks like a duck and quacks like a duck…”. Learn Motiv 36, 190–207, doi:10.1016/j.lmot.2005.02.006 (2005).

8 Eichenbaum, H., Sauvage, M., Fortin, N., Komorowski, R. & Lipton, P. Towards a functional organization of episodic memory in the medial temporal lobe. Neurosci Biobehav R 36, 1597–1608, doi:10.1016/j.neubiorev.2011.07.006 (2012).

9 Burgess, N., Maguire, E. A. & O’Keefe, J. The human hippocampus and spatial and episodic memory. Neuron 35, 625–641, doi:Doi 10.1016/S0896-6273(02)00830-9 (2002).

10 Cabeza, R., Ciaramelli, E., Olson, I. R. & Moscovitch, M. The parietal cortex and episodic memory: an attentional account. Nat Rev Neurosci 9, 613–625, doi:10.1038/nrn2459 (2008).

11 Sestieri, C., Shulman, G. L. & Corbetta, M. The contribution of the human posterior parietal cortex to episodic memory. Nat Rev Neurosci 18, 183–192, doi:10.1038/nrn.2017.6 (2017).

12 Healy, A. F. Coding of temporal-spatial patterns in short-term memory. Oct 1975. Journal of Verbal Learning & Verbal Behavior. 14, pp, doi:10.1016/S0022-5371%2875%2980026-0 (1975).

13 Rondina, R., 2nd, Curtiss, K., Meltzer, J. A., Barense, M. D. & Ryan, J. D. The organisation of spatial and temporal relations in memory. Memory 25, 436–449, doi:10.1080/09658211.2016.1182553 (2017).

14 van Asselen, M., Van der Lubbe, R. H. & Postma, A. Are space and time automatically integrated in episodic memory? Memory 14, 232–240, doi:10.1080/09658210500172839 (2006).

15 Delogu, F., Nijboer, T. C. & Postma, A. Encoding location and serial order in auditory working memory: evidence for separable processes. Cognitive processing 13, 267–276, doi:10.1007/s10339-012-0442-3 (2012).

16 Delogu, F., Nijboer, T. C. & Postma, A. Binding “When” and “Where” Impairs Temporal, but not Spatial Recall in Auditory and Visual Working Memory. Front Psychol 3, 62, doi:10.3389/fpsyg.2012.00062 (2012).

17 Dutta, A. & Nairne, J. S. The separability of space and time: dimensional interaction in the memory trace. Mem Cognit 21, 440–448 (1993).

18 Healy, A. F. Separating item from order information in short-term memory. Dec 1974. Journal of Verbal Learning & Verbal Behavior. 13, pp, doi:10.1016/S0022-5371%2874%2980052-6 (1974).

19 Kohler, S., Moscovitch, M. & Melo, B. Episodic memory for object location versus episodic memory for object identity: do they rely on distinct encoding processes? Mem Cognit 29, 948–959 (2001).

20 Wittig, J. H., Jr., Morgan, B., Masseau, E. & Richmond, B. J. Humans and monkeys use different strategies to solve the same short-term memory tasks. Learn Mem 23, 644–647, doi:10.1101/lm.041764.116 (2016).

21 Sakamoto, Y. & Love, B. C. Vancouver, Toronto, Montreal, Austin: Enhanced oddball memory through differentiation, not isolation. Psychon B Rev 13, 474–479, doi:Doi 10.3758/Bf03193872 (2006).

22 Ebbinghaus, H. Memory: a contribution to experimental psychology. Annals of neurosciences 20, 155–156, doi:10.5214/ans.0972.7531.200408 (2013).

23 Boran, E. et al. Persistent hippocampal neural firing and hippocampal-cortical coupling predict verbal working memory load. Sci Adv 5, eaav3687, doi:10.1126/sciadv.aav3687 (2019).

24 Jeneson, A., Mauldin, K. N., Hopkins, R. O. & Squire, L. R. The role of the hippocampus in retaining relational information across short delays: the importance of memory load. Learn Mem 18, 301–305, doi:10.1101/lm.2010711 (2011).

25 Kaminski, J. et al. Persistently active neurons in human medial frontal and medial temporal lobe support working memory. Nat Neurosci 20, 590–601, doi:10.1038/nn.4509 (2017).

26 Conrad, R. Order error in immediate recall of sequences. 1965. Journal of Verbal Learning & Verbal Behavior. 4, pp, doi:10.1016/S0022-5371%2865%2980015-9 (1965).

27 Murdock, B. B., Jr. & Vom Saal, W. Transpositions in short-term memory. Journal of Experimental Psychology. 74, pp, doi:10.1037/h0024507 6032566 (1967).

28 Estes, W. K. Processes of memory loss, recovery, and distortion. Psychol Rev 104, 148–169, doi:10.1037/0033-295x.104.1.148 (1997).

29 Nairne, J. S. N. I; Serra, M; Byun, E. Positional distinctiveness and ratio rule in free recall. J Mem Lang 37, 155–166 (1997).

30 Marshall, P. H. & Werder, P. R. The effects of the elimination of rehearsal on primacy and recency. Journal of Verbal Learning and Verbal Behavior 11, 649–653, doi: https://doi.org/10.1016/S0022-5371(72)80049-5 (1972).

31 Murdock Jr., B. B. Effects of a subsidiary task on short-term memory. British Journal of Psychology 56, 413–419, doi:10.1111/j.2044-8295.1965.tb00983.x (1965).

32 Alvarez, G. A. & Cavanagh, P. The capacity of visual short-term memory is set both by visual information load and by number of objects. Psychol Sci 15, 106–111, doi:10.1111/j.0963-7214.2004.01502006.x (2004).

33 Luck, S. J. & Vogel, E. K. The capacity of visual working memory for features and conjunctions. Nature 390, 279–281, doi:10.1038/36846 (1997).

34 van Vugt, M. K., Schulze-Bonhage, A., Litt, B., Brandt, A. & Kahana, M. J. Hippocampal gamma oscillations increase with memory load. J Neurosci 30, 2694–2699, doi:10.1523/JNEUROSCI.0567-09.2010 (2010).

35 Schacter, D. L., Norman, K. A. & Koutstaal, W. The cognitive neuroscience of constructive memory. Annu Rev Psychol 49, 289–318, doi:10.1146/annurev.psych.49.1.289 (1998).

36 Brainard, D. H. The psychophysics toolbox. Spatial Vision 10, 433–436, doi:Doi 10.1163/156856897x00357 (1997).

37 Meeker, W. Q. & Escobar, L. A. Statistical methods for reliability data. (Wiley, 1998).

38 Huelsenbeck, J. P. & Crandall, K. A. Phylogeny estimation and hypothesis testing using maximum likelihood. Annu Rev Ecol Syst 28, 437–466, doi:Doi 10.1146/Annurev.Ecolsys.28.1.437 (1997).

